# Time-of-day related fluctuations of self-belief formation

**DOI:** 10.1101/2024.12.05.627011

**Authors:** Annalina V. Mayer, Alexander Schröder, Nora Czekalla, Laura Müller-Pinzler, Laura Rosenbusch, Frieder M. Paulus, Henrik Oster, Clara Sayk, Mathias Kammerer, Ines Wilhelm-Groch, Sören Krach

## Abstract

Time of day influences a variety of human cognitive processes, including attention, executive functions and memory formation, as well as affective experiences and mood. However, circadian modulations of self-related learning and belief formation, which are highly affected by emotional states, remain poorly understood.

Here, we present results from exploratory post-hoc analyses on data aggregated from five studies assessing the formation of self-related ability beliefs. A total of *N*=242 healthy participants completed a validated learning task at different times of the day, during which they continuously received feedback on their performance. Computational modeling was applied to quantify participants’ learning behavior during the task.

Results suggest an association between time-of-day and self-belief formation, showing that participants who were tested in the evening (7:00-9:59 p.m.) updated their self-beliefs more strongly in response to the received feedback compared to those tested in the afternoon (1:00-3:59 p.m.). Evidence from additional models indicated that these differences were driven by non-linear, rhythmic changes in self-belief formation across different times of the day.

Future studies should systematically examine within-subject fluctuations in self-belief formation across the day and address the influence of individual factors such as chronotype, age, mood and sleep quality. Understanding circadian modulations of self-related belief formation could contribute to optimized interventions for conditions characterized by maladaptive self-beliefs, such as depression, as well as in academic contexts.

## Introduction

Time of day influences a variety of human cognitive functions [1,2]. For example, different components of attention, including tonic alertness, phasic alertness and selective attention, have been found to vary according to the time of day [3]. Performance in these areas generally declines during nighttime and early morning hours, improves around noon, and peaks in the afternoon and evening [4]. Similar results have been found for memory [5,6] and executive functions, including inhibition, flexibility, and self-monitoring [7–9]. These cognitive functions also generally improve during the day and decline at night and in the early morning. Circadian rhythms in attention, executive functions and memory can impact higher-order cognitive functions such as decision-making and problem-solving [10]. This could have important implications for work environments, as the time-of-day related decline in these cognitive functions may increase the risk of errors and accidents during night shifts and early morning commutes [8,9]. Additionally, individual differences in circadian typology (i.e., morning vs. evening chronotype) can influence cognitive performance, with some tasks showing optimal results when individuals are tested at their preferred time of day [11,12].

While circadian modulations of attention, executive functions and memory have been well studied, less is known about time-of-day effects on learning in humans. The variety of tasks and measures used in this area of research make it difficult to compare results and draw conclusions for learning in general. For sequence learning, it has been shown that time of day affects the expression of learned skills, with task performance decreasing in the evening [13]. Using an associative learning task, older adults were found to have greater intraindividual variability in the evening, but not young adults [14]. An analysis of panel data suggested that students learned more in the morning than later in the school day, which was reflected in higher test scores when classes were held in the morning [15]. Taken together, there is evidence that time of day can influence learning in humans, although the specific mechanisms remain to be understood.

In addition to cognitive functioning, the time of day influences emotional experience and mood [16]. Positive affect exhibits a 24-hour temporal variation, following a sinusoidal pattern akin to the circadian rhythm of body temperature [17]. Diary studies revealed that participants reported more positive and fewer negative emotions in the evening [18] with a peak of positive affect in the afternoon [19]. The 24-hour mood variation is also reflected in the content of social media posts [20]. Among adolescents, mood has been observed to improve progressively throughout the school day [21]. Additionally, emotional reactivity and emotion regulation are subject to circadian regulation [22] and circadian rhythm disturbances play an important role in the pathophysiology of mood disorders [23].

To date, no study has addressed time-of-day effects in self-related learning or self-belief formation. Self-belief formation refers to the process of integrating information from the environment to arrive at a belief about “who one is” [24]. This process can be formalized using computational models derived from reinforcement learning [25,26]. In these models, self-belief formation is described as a series of belief updates that are inherently connected [24]. Beliefs about the self, for example about one’s appearance, abilities or popularity, change in response to prediction errors, that is, the mismatch between expected and actual feedback. A model-specific learning rate determines the degree to which prediction errors are used to update these self-beliefs. High learning rates indicate stronger adaptation to changing feedback, and low learning rates indicate weaker adaptation [25].

Learning rates can vary between different learning contexts and are often influenced by affective states and motives [24,27]. For example, when individuals strive for a positive self-image, the process of self-belief formation is often positively biased [28]. In this case, this means that self-beliefs are updated more strongly after better-than-expected feedback (i.e., higher learning rate) compared to worse-than-expected feedback (i.e., lower learning rate) [29]. Similarly, it has been shown that when people are in a low mood, they tend to update their self-beliefs less positively [30]. As both mood and cognitive functions are subject to diurnal fluctuations, the formation of self-beliefs could also vary depending on the time of day.

To explore this possibility, we aggregated data from five previous studies that examined self-belief formation in healthy participants. All studies involved a validated self-related learning task that participants completed at different times of the day. Computational modeling was applied to quantify participants’ learning behavior. We then explored time-of-day effects on individual learning rates on a between-subject level.

## Methods

### Participants

This exploratory post-hoc analysis included data of a total of *N*=242 participants (*n*=176 female) from five previous studies assessing self-belief formation (unpublished and published data [26,27,31,32]). The final sample comprised only healthy adults between the ages of 18 and 35 (*M* = 23.6, *SD* = 3.57) without any psychiatric or neurological disorders. All participants were fluent in German, had normal or corrected-to-normal vision, and provided written consent. All five original studies adhered to the ethical standards of the American Psychological Association and were approved by the ethics committee of the University of Lübeck.

### Learning of Own Performance Task (LOOP)

In all studies, participants completed the Learning of Own Performance Task (LOOP), a learning task that uses trial-by-trial feedback in a performance context which allows the formation of novel self-beliefs. In each trial, individuals received feedback about their ability to estimate various properties or characteristics like the weight of different animals or the height of buildings (**Figure 1A**). This task was validated in multiple behavioral and neuroimaging studies [26,27,31–33].

**Figure 1.**
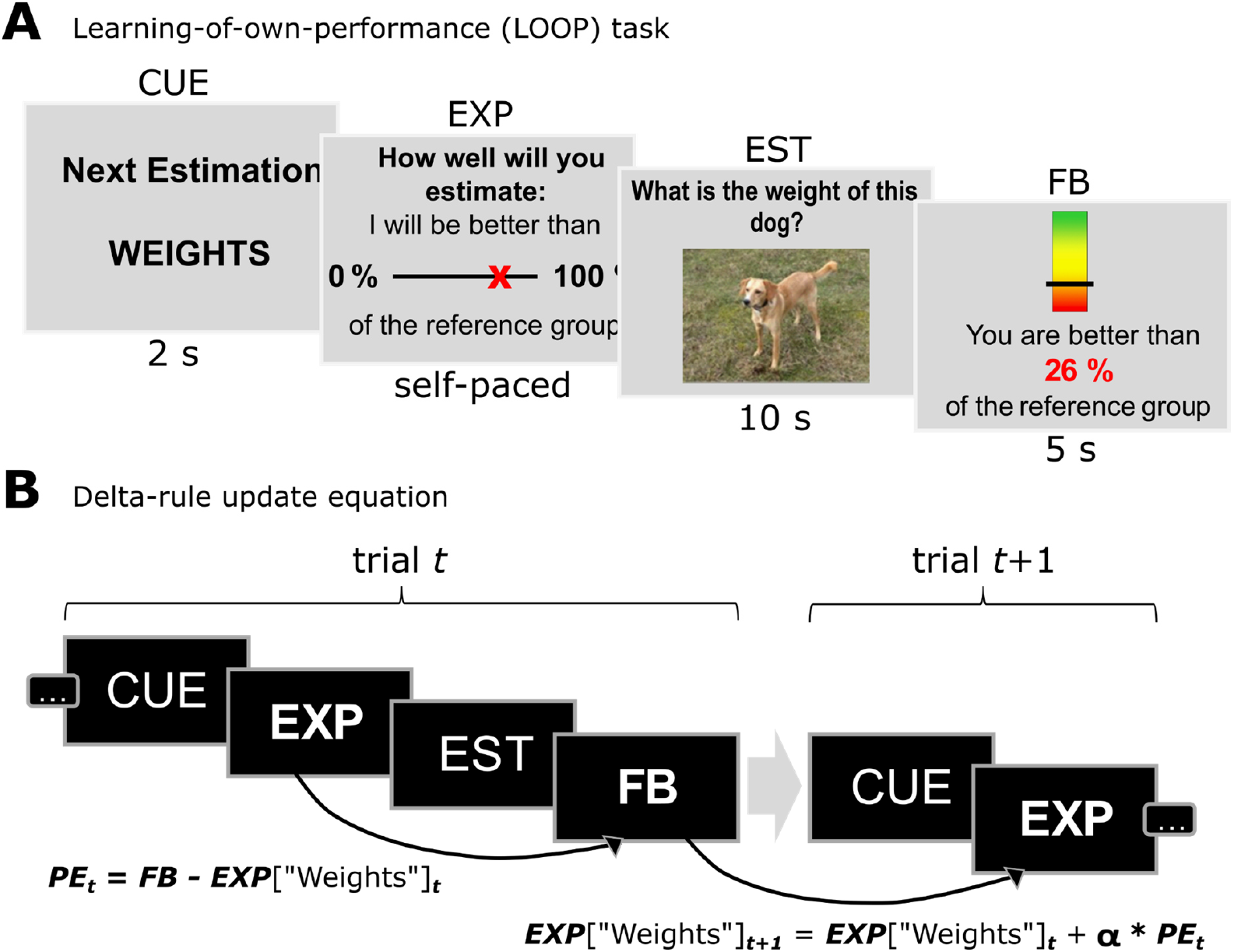
Learning-of-own-performance (LOOP) task. **a**. Each trial began with the presentation of a category cue (CUE). Participants were then prompted to provide a performance expectation (EXP). This was followed by an estimation question (EST). At the end of each trial, feedback (FB) was provided. **b**. Delta-rule update equation based on the Rescorla-Wagner model. A prediction error (PE) was computed from the difference between the performance expectation (EXP) and the feedback (FB) of the corresponding trial. The expectation for the next trial within the same category was updated by incorporating the prior prediction error. This update was modulated by a learning rate (α), which determined the extent to which the prediction error influenced adjustments to performance expectations.

There were two categories of estimation tasks. Unbeknownst to the participants, the presented feedback followed a fixed sequence, with one category (e.g., the height of buildings) being associated with more positive feedback (high ability condition: positive prediction errors in 70% of all trials), and the other category (e.g., the weight of animals) being associated with more negative feedback (low ability condition: positive prediction errors in 30% of all trials) (**Figure 2A**). Each trial started with a cueing phase indicating the estimation category of the respective trial. Participants were then asked to rate their expected performance (EXP) for the upcoming task by ranking themselves on a percentile scale relative to a supposed reference group of previous participants. Next, the estimation question followed, during which participants had ten seconds to answer on a continuous scale of plausible options. After this, performance feedback (FB) was provided on the same scale as the previous expectation rating (e.g., “You are better than 65% of the reference group.”) for three seconds. To maintain variability in feedback and to keep prediction errors independent of participants’ expectations, trial-wise feedback was determined by a sequence of fixed prediction errors. The entire task consisted of 20 trials per ability condition. After task completion, participants were debriefed about the manipulated feedback and compensated for their participation.

**Figure 2.**
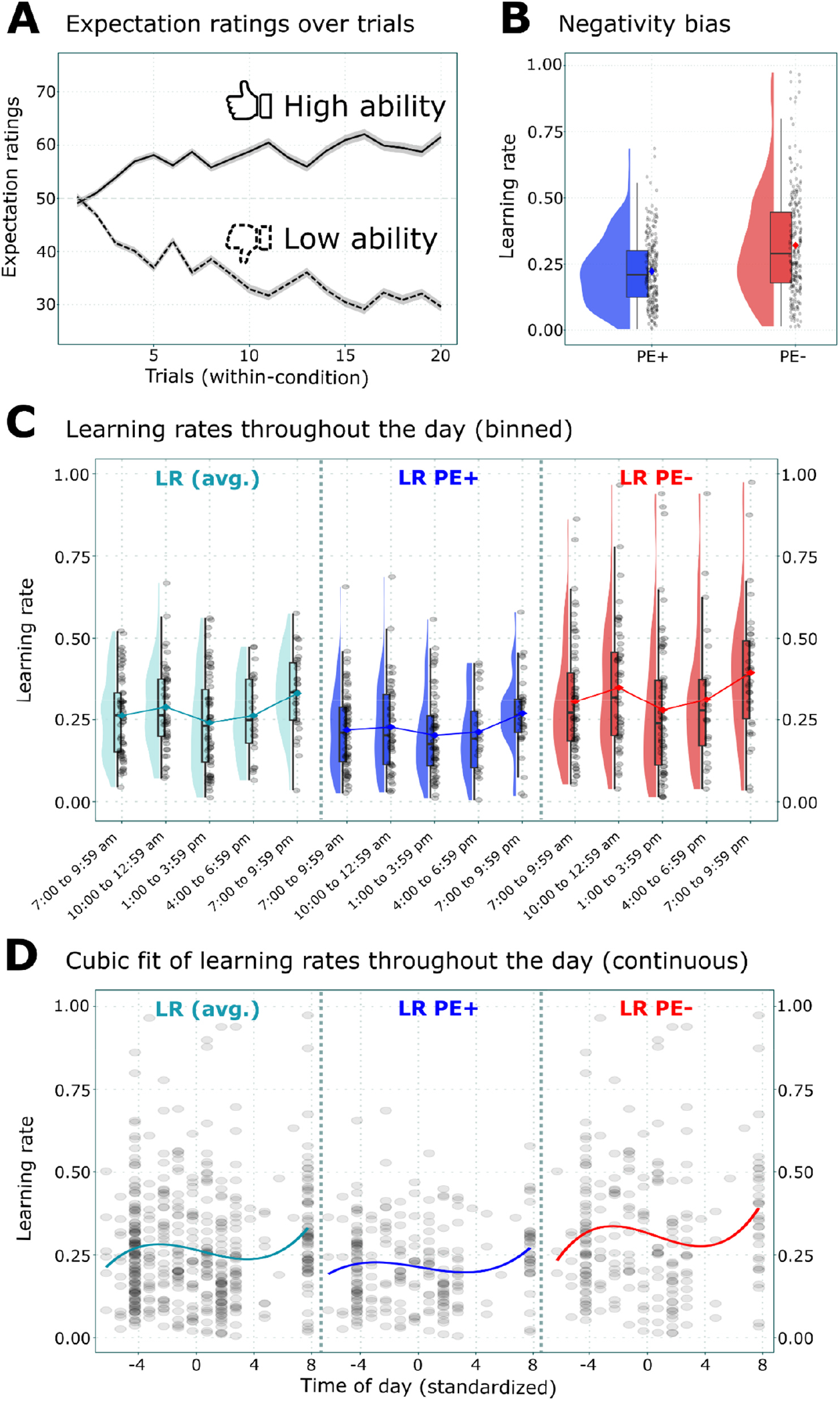
Fluctuations of self-related learning over the day. **a**. Expectation ratings over the trials of the LOOP-task. Participants adjusted their expectations depending on the feedback condition. **b**. Self-related negativity bias. Participants exhibited significantly higher learning rates for updates after negative prediction errors. **c**. Learning rates (separated by PE valence) throughout the day (binned). **d**. Cubic fit of learning rates (separated by PE valence) throughout the day.

### Computational modeling

Trial-by-trial self-belief changes were quantified using Rescorla-Wagner-based reinforcement learning equations [25,34] (**Figure 1B**). The basic learning model reads as follows (1):

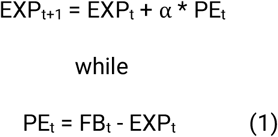

“EXP” denotes the performance expectation rating, “FB” stands for the received feedback, “t” indexes the within-category trial, “PE” represents the prediction error, and “α” is the learning rate. The learning rate represents a weight on the prediction error. Its height determines how much of the experienced deviation between expectation and feedback is used to update one’s expectation. The best-performing model of the most recent studies that used the LOOP task extended this base model in multiple ways [27,31,32]: First, different learning rates for updates following positive vs. negative PEs were introduced. Second, a weighting parameter *w* was added that reduces the impact of the learning rates when the feedback approaches the extremes of the scale (near 0% or 100% percentiles). This additional parameter *w* is multiplied by the relative probability density of the normal distribution (ND) associated with each possible feedback percentile value. This addition to the base model is based on the assumption that more extreme feedback is perceived to be less probable and indicative. The final model (2) reads as follows, where “valence” depicts the valence of the received PE, i.e. whether a PE was positive or negative (2):

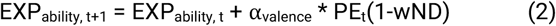

For the analyses of this manuscript, all available data were re-estimated using this model.

Model fitting was conducted using the *RStan* package [35] for R [36]. For all participants, the model was fitted individually by employing Markov Chain Monte Carlo (MCMC) sampling. In total, 3,400 samples were drawn (thereof 1,000 burn-in samples), thinned by a factor of three. The convergence of the Markov chains was checked visually, and the *Rhat* values for each parameter were inspected. Effective sample sizes *n*_*ef*f_were checked to be mostly bigger than 1,500. These values describe the effective number of independent draws from the posterior distributions of the model’s parameters. After successful model estimation, the model parameters like learning rates were summarized by calculating the mean values of the posterior distributions per participant. These summarized estimates were used in the subsequent analyses.

### Statistical analyses

All statistical analyses were conducted using the R software for statistical computations [36]. Exploratory post-hoc analyses started with a visual inspection of descriptive data on learning rates from the computational modeling plotted over the time of day.

To examine whether the time of day of participation influenced self-belief formation, learning rates were analyzed using Linear Mixed-Effects Models (LMMs) implemented via the *lme4* package in R [37]. Data were categorized by time-of-day intervals (factor: time) as follows: 7:00 to 9:59 am, 10:00 to 12:59 am, 1:00 to 3:59 pm, 4:00 to 6:59 pm, and 7:00 to 9:59 pm. The maximal LMM included the between-subject factor time, the within-subject factor PE-valence (learning rate α_PE+_vs. α_PE-_), the interaction term time × PE-valence, and the intercept as fixed effects. Subject-level intercepts were included as random effects. Model construction began with a baseline model containing only the intercept as a fixed effect, with additional fixed effects added iteratively. Model estimation was employed at this stage using Maximum Likelihood (ML) since the models only differed in the fixed-effects structure. Likelihood ratio tests determined the model that best explained the data. Subsequently, the winning model was re-estimated using Restricted Maximum Likelihood (REML) with significance tests for estimates using Satterthwaite’s approximation for degrees of freedom [38]. To explore specific main effects and interactions, pairwise comparisons of estimated marginal means were calculated using the package *emmeans* [39].

Further tests were conducted to identify potential non-linear effects of time of day. For this, polynomial LMMs were used to test linear, quadratic, and cubic fits. Learning rates as dependent variables were considered in separate models with the standardized time of day as a continuous predictor. The model construction procedure was analogous to the one previously explained. In this case, we began with a linear baseline model that included the intercept, standardized time of day, and PE valence as fixed effects. Subject-level intercepts were included as random effects. From this, polynomial predictors and interaction terms were added iteratively. Again, the model fitting the data best was subsequently re-estimated using REML and served for further analyses.

## Results

### Learning rates differ across the day

The model containing only main effects of time and PE valence, but no interaction effect, best explained the data (R^2^ marginal = 0.11, R^2^ conditional = 0.33). As reported in previous publications [26,27,31–33], there was a significant effect of PE valence on self-belief formation (learning rates, β = 0.1, *p* < .001): Learning rates were significantly higher for negative PEs, indicating that participants updated their performance expectations more strongly following worse-than-expected feedback (**Figure 2B**). Notably, there was also a significant effect of time of day on learning rates. Pairwise comparisons of time intervals revealed that learning rates were higher in the evening (7:00 to 9:59 pm) compared to the early afternoon (1:00 to 3:59 pm, *t*(237) = -3.31, *p* < .01, **Figure 2C**), suggesting overall stronger self-belief updating during this time. Other comparisons did not reach significance when controlling for multiple testing (see **Table 1** for all pairwise comparisons).

**Table 1.** Pairwise comparisons of estimated marginal means of learning rates across time-of-day intervals.

| contrast | estimate | SE | df | t | $p_{uncorrected}$ | $p_{Tukey}$ |
| --- | --- | --- | --- | --- | --- | --- |
| 7:00-9:59am - 10:00-10:59am | -0.03 | 0.02 | 237 | -1.06 | .29 | .826 |
| 7:00-9:59am - 1:00-3:00pm | 0.02 | 0.02 | 237 | 0.96 | .339 | .874 |
| 7:00-9:59am - 4:00-6:59pm | 0.00 | 0.03 | 237 | 0.01 | .995 | 1 |
| 7:00-9:59am - 7:00-9:59pm | -0.07 | 0.03 | 237 | -2.52 | .012* | .089 |
| 10:00-12:59am - 1:00-3:00pm | 0.05 | 0.02 | 237 | 1.96 | .052 | .291 |
| 10:00-12:59am - 4:00-6:59pm | 0.03 | 0.03 | 237 | 0.82 | .415 | .926 |
| 10:00-12:59am - 7:00-9:59pm | -0.04 | 0.03 | 237 | -1.51 | .132 | .555 |
| 1:00-3:00pm - 4:00-6:59pm | -0.02 | 0.03 | 237 | -0.70 | .482 | .955 |
| 1:00-3:00pm - 7:00-9:59pm | -0.09 | 0.03 | 237 | -3.31 | .001*** | .009** |
| 4:00-6:59pm - 7:00-9:59pm | -0.07 | 0.03 | 237 | -2.03 | .044* | .256 |

Note. SE = standard error of the mean; df = degrees of freedom. \* $p < .05$ ; \*\* $p < .01$ ; \*\*\* $p \leq .001$ . $p$ -values are corrected for multiple comparisons using Tukey's approach.

### Cubic fluctuations in time-of-day effects on learning rates

Linear and polynomial LMMs were used to explore the possibility of non-linear rhythmic fluctuations of learning rates. The model that best explained the data included the main effect of PE valence and linear, quadratic, and cubic terms for time of day, but no interaction effect of time of day and PE valence (R^2^ marginal = 0.1; R^2^ conditional = 0.33). Again, there was a main effect of PE valence suggesting an overall negativity bias (β = 0.1, 95% CI [0.07, 0.12], *p* < .001). More notably, there was a significant main effect for a cubic fit of time of day (β = 3.99e-4, 95% CI [3.38e-5, 7.63e-4], *p* < .05). This supports the idea of non-linear rhythmic fluctuations in belief formation depending on the time of day. Visual inspection of learning rates by valence (**Figure 2C**) as well as cubic fit plots (**Figure 2D**) suggest that these rhythmic variations may be driven particularly by learning rates following negative prediction errors, although the winning models did not include an interaction term of time of day and PE valence.

## Discussion

Using data from five previous studies, we show that self-belief formation fluctuates across different times of the day, with a decline in learning rates in the afternoon and an increase during the evening hours. In other words, participants who were tested in the evening updated their self-beliefs more strongly in response to the received feedback compared to those tested in the afternoon. Our results suggest that these differences are driven by non-linear, rhythmic changes in learning across different times of the day. The pattern of fluctuations partly resembles those of other cognitive functions, which show a decline during night and early morning and peak in the evening [1]. The decline of self-related learning rates in the afternoon might be related to a so-called post-noon dip in vigilance [40]. Vigilance refers to the ability to detect and react to stimuli in the environment over prolonged periods of time [41]. A decline in vigilance may thus also lead to a reduced receptiveness to self-related information and thus influence the formation of self-beliefs. As reported in previous work, we also demonstrate that learning about one’s own abilities is overall negatively biased, with significantly higher learning rates for worse-than-expected compared to better-than-expected feedback [26].

Diurnal variations in self-belief formation, along with general fluctuations in cognitive functioning, suggest that people may be more receptive to self-related feedback at certain times of the day. This might have important implications for any context where individuals form new beliefs about themselves. In psychotherapy, for example, one central goal is often to revise maladaptive negative self-beliefs [42,43], which are seen as a key factor in the onset and maintenance of various mental disorders [44]. Further, self-beliefs shape decision-making [45] and are related to academic achievement, with studies showing that those with more positive beliefs about their abilities actually perform better in academic contexts [46,47]. Understanding how self-related information processing and self-belief formation change throughout the day could help identify the best times for interventions in schools and therapy. This, in turn, could support the development of adaptive self-beliefs, greater self-efficacy, and improved academic performance.

To reach this goal, further studies are needed that consider the following aspects. First, studies should systematically test the same subjects at different times of the day to quantify fluctuations on a within-subject level. Second, there are several individual factors that could potentially influence the course of self-belief formation across the day that should be addressed in future studies. One such factor might be *chronotype*, which reflects a person’s preferred period of alertness and energy during the sleep-wake cycle [48]. Chronotype can indicate an optimal time to perform feedback learning tasks in general [11] and may similarly interact with learning from self-related feedback. Since chronotype changes over the lifespan [49], *age* is an also important factor to include when studying diurnal variations of self-belief formation. Diurnal variations in *mood* are another critical consideration, as they exhibit strong individual differences partially influenced by chronotype [50]. To better understand self-specific belief formation, which is closely tied to affective states [27], future studies should investigate how mood interacts with updating self-beliefs at different times of the day. Additionally, *sleep quality* is closely linked to both affective states [51] and chronotype, together with the degree of alignment between an individual’s biological rhythms and societal schedules—a phenomenon known as “social jetlag” [52]. To account for these interactions, sleep quality should be recorded alongside chronotype in future research.

Together, this study highlights diurnal fluctuations in self-belief formation, with evidence suggesting diminished updates during the early afternoon. These findings extend the literature on circadian variations in cognitive and affective processes with a special focus on self-related cognitions, underscoring the importance of considering time-of-day effects in understanding belief formation. Future research should investigate within-subject fluctuations as well as individual differences to deepen our understanding of how temporal and individual factors interact in self-related learning. These insights could inform interventions aimed at promoting adaptive self-beliefs in therapeutic settings or identify optimal learning windows in school or work contexts to enhance self-efficacy beliefs.

## Notes

### Competing Interest Statement

The authors have declared no competing interest.

## References

1. Schmidt C, Collette F, Cajochen C, Peigneux P. A time to think: circadian rhythms in human cognition. Cogn Neuropsychol. 2007;24: 755–789. doi:10.1080/02643290701754158

2. Salehinejad MA, Wischnewski M, Ghanavati E, Mosayebi-Samani M, Kuo M-F, Nitsche MA. Cognitive functions and underlying parameters of human brain physiology are associated with chronotype. Nat Commun. 2021;12: 4672. doi:10.1038/s41467-021-24885-0

3. Valdez P, Ramírez C, García A, Talamantes J, Armijo P, Borrani J. Circadian rhythms in components of attention. Biol Rhythm Res. 2005;36: 57–65. doi:10.1080/09291010400028633

4. Valdez P. Circadian rhythms in attention. Yale J Biol Med. 2019;92: 81–92. Available: https://www.ncbi.nlm.nih.gov/pmc/articles/PMC6430172/

5. Folkard S, Monk TH. Circadian rhythms in human memory. Br J Psychol. 1980;71: 295–307. doi:10.1111/j.2044-8295.1980.tb01746.x

6. Potter D, Keeling D. Effects of moderate exercise and circadian rhythms on human memory. Journal of Sport and Exercise Psychology. 2005;27: 117–125. Available: https://journals.humankinetics.com/view/journals/jsep/27/1/article-p117.xml

7. Ramírez C, García A, Valdez P. Identification of circadian rhythms in cognitive inhibition and flexibility using a Stroop task: Circadian rhythms in executive functions. Sleep Biol Rhythms. 2012;10: 136–144. doi:10.1111/j.1479-8425.2012.00540.x

8. García A, Ramírez C, Valdez P. Circadian variations in self-monitoring, a component of executive functions. Biol Rhythm Res. 2016;47: 7–23. doi:10.1080/09291016.2015.1075722

9. García A, Ramírez C, Martínez B, Valdez P. Circadian rhythms in two components of executive functions: cognitive inhibition and flexibility. Biol Rhythm Res. 2012;43: 49–63. doi:10.1080/09291016.2011.638137

10. Blatter K, Cajochen C. Circadian rhythms in cognitive performance: methodological constraints, protocols, theoretical underpinnings. Physiol Behav. 2007;90: 196–208. doi:10.1016/j.physbeh.2006.09.009

11. Bennett CL, Petros TV, Johnson M, Ferraro FR. Individual differences in the influence of time of day on executive functions. The American journal of psychology. 2008;121: 349–361. Available: https://scholarlypublishingcollective.org/uip/ajp/article-abstract/121/3/349/258607

12. Lara T, Madrid JA, Correa Á. The vigilance decrement in executive function is attenuated when individual chronotypes perform at their optimal time of day. PLoS One. 2014;9: e88820. doi:10.1371/journal.pone.0088820

13. Keisler A, Ashe J, Willingham D. Time of day accounts for overnight improvement in sequence learning. Learn Mem. 2007;14: 669–672. doi:10.1101/LM.751807

14. Hogan MJ, Kelly CAM, Verrier D, Newell J, Hasher L, Robertson IH. Optimal time-of-day and consolidation of learning in younger and older adults. Exp Aging Res. 2009;35: 107–128. doi:10.1080/03610730802545366

15. Pope N. How the time of day affects productivity: Evidence from school schedules. Rev Econ Stat. 2016;98: 1–11. doi:10.1162/REST_a_00525

16. Watson D. Mood and Temperament. New York, NY: Guilford Publications; 2000. Available: https://books.google.com/books/about/Mood_and_Temperament.html?id=iPFboulhcQcC

17. Murray G, Allen NB, Trinder J. Mood and the circadian system: investigation of a circadian component in positive affect. Chronobiol Int. 2002;19: 1151–1169. doi:10.1081/cbi-120015956

18. English T, Carstensen LL. Emotional experience in the mornings and the evenings: consideration of age differences in specific emotions by time of day. Front Psychol. 2014;5: 185. doi:10.3389/fpsyg.2014.00185

19. Egloff B, Tausch A, Kohlmann C-W, Krohne HW. Relationships between time of day, day of the week, and positive mood: Exploring the role of the mood measure. Motiv Emot. 1995;19: 99–110. doi:10.1007/bf02250565

20. Dzogang F, Lightman S, Cristianini N. Circadian mood variations in Twitter content. Brain Neurosci Adv. 2017;1: 2398212817744501. doi:10.1177/2398212817744501

21. Díaz-Morales JF, Escribano C, Jankowski KS. Chronotype and time-of-day effects on mood during school day. Chronobiol Int. 2015;32: 37–42. doi:10.3109/07420528.2014.949736

22. Tucker AM, Feuerstein R, Mende-Siedlecki P, Ochsner KN, Stern Y. Double dissociation: circadian off-peak times increase emotional reactivity; aging impairs emotion regulation via reappraisal. Emotion. 2012;12: 869–874. doi:10.1037/a0028207

23. Germain A, Kupfer DJ. Circadian rhythm disturbances in depression. Hum Psychopharmacol. 2008;23: 571–585. doi:10.1002/hup.964

24. Krach S, Müller-Pinzler L, Czekalla N, Schröder A, Wilhelm-Groch I, Luebber F, et al. Examining self-belief formation through artificial beliefs. 2024. Available: https://www.researchgate.net/profile/Soeren-Krach-2/publication/380792680_Examining_self-belief_formation_through_artificial_beliefs/links/664f1d0dbc86444c72f9e2b5/Examining-self-belief-formation-through-artificial-beliefs.pdf

25. Zhang L, Lengersdorff L, Mikus N, Gläscher J, Lamm C. Using reinforcement learning models in social neuroscience: frameworks, pitfalls and suggestions of best practices. Soc Cogn Affect Neurosci. 2020;15: 695–707. doi:10.1093/scan/nsaa089

26. Müller-Pinzler L, Czekalla N, Mayer AV, Stolz DS, Gazzola V, Keysers C, et al. Negativity-bias in forming beliefs about own abilities. Sci Rep. 2019;9: 14416. doi:10.1038/s41598-019-50821-w

27. Müller-Pinzler L, Czekalla N, Mayer AV, Schröder A, Stolz DS, Paulus FM, et al. Neurocomputational mechanisms of affected beliefs. Communications Biology. 2022;5: 1–15. doi:10.1038/s42003-022-04165-3

28. Eil D, Rao JM. The good news-bad news effect: Asymmetric processing of objective information about yourself. Am Econ J Microecon. 2011;3: 114–138. doi:10.1257/mic.3.2.114

29. Sharot T, Garrett N. Forming Beliefs: Why Valence Matters. Trends Cogn Sci. 2016;20: 25–33. doi:10.1016/j.tics.2015.11.002

30. Kube T, Korn C. Induced negative affect hinders self-referential belief updating in response to social feedback. Emotion. 2024. doi:10.1037/emo0001426

31. Czekalla N, Schröder A, Mayer AV, Stierand J, Stolz DS, Kube T, et al. Neurocomputational mechanisms underlying maladaptive self-belief formation in depression. bioRxiv. 2024. p. 2024.05.09.593087. doi:10.1101/2024.05.09.593087

32. Schröder A, Czekalla N, Mayer AV, Zhang L, Stolz DS, Korn CW, et al. Prior expectations about own abilities bias self-belief formation and hinder subsequent revision. bioRxiv. 2024. p. 2024.08.30.610443. doi:10.1101/2024.08.30.610443

33. Czekalla N, Stierand J, Stolz DS, Mayer AV, Voges JF, Rademacher L, et al. Self-beneficial belief updating as a coping mechanism for stress-induced negative affect. Sci Rep. 2021;11: 17096. doi:10.1038/s41598-021-96264-0

34. Rescorla R, Wagner A. A theory of Pavlovian conditioning: The effectiveness of reinforcement and non-reinforcement. Classical Conditioning: Current Research and Theory. 1972.

35. Stan Development Team. RStan: the R interface to Stan. 2021. Available: https://mc-stan.org/

36. R Core Team. R: A Language and Environment for Statistical Computing. Vienna, Austria: R Foundation for Statistical Computing; 2021. Available: https://www.R-project.org/

37. Bates D, Mächler M, Bolker B, Walker S. Fitting Linear Mixed-Effects Models Using lme4. Journal of Statistical Software. 2015. pp. 1–48. doi:10.18637/jss.v067.i01

38. Meteyard L, Davies RAI. Best practice guidance for linear mixed-effects models in psychological science. J Mem Lang. 2020;112: 104092. doi:10.1016/j.jml.2020.104092

39. Lenth RV. emmeans: Estimated Marginal Means, aka Least-Squares Means. 2023. Available: https://CRAN.R-project.org/package=emmeans

40. Valdez P, Ramírez C, García A. Circadian rhythms in cognitive performance: implications for neuropsychological assessment. ChronoPhysiology and Therapy. 2012 [cited 28 Nov 2024]. doi:10.2147/CPT.S32586

41. Frankmann JP, Adams JA. Theories of vigilance. Psychological Bulletin. 1962;59: 257–272. doi:10.1037/H0046142

42. Beck AT. Thinking and depression. Ii. Theory and therapy. Arch Gen Psychiatry. 1964;10: 561–571. doi:10.1001/ARCHPSYC.1964.01720240015003

43. Beck AT. Thinking and depression. I. Idiosyncratic content and cognitive distortions. Arch Gen Psychiatry. 1963;9: 324–333. doi:10.1001/ARCHPSYC.1963.01720160014002

44. Rief W, Glombiewski JA, Gollwitzer M, Schubö A, Schwarting R, Thorwart A. Expectancies as core features of mental disorders. Curr Opin Psychiatry. 2015;28: 378–385. doi:10.1097/YCO.0000000000000184

45. Krueger N, Dickson PR. How believing in ourselves increases risk taking: Perceived self-efficacy and opportunity recognition. Decis Sci. 1994;25: 385–400. doi:10.1111/j.1540-5915.1994.tb00810.x

46. Valentine JC, DuBois DL, Cooper H. The relation between self-beliefs and academic achievement: A meta-analytic review. Educ Psychol. 2004;39: 111–133. doi:10.1207/s15326985ep3902_3

47. Bandura A, Barbaranelli C, Caprara GV, Pastorelli C. Multifaceted impact of self-efficacy beliefs on academic functioning. Child Dev. 1996;67: 1206–1222. doi:10.1111/j.1467-8624.1996.tb01791.x

48. Roenneberg T. Having trouble typing? What on earth is chronotype? J Biol Rhythms. 2015;30: 487–491. doi:10.1177/0748730415603835

49. Fischer D, Lombardi DA, Marucci-Wellman H, Roenneberg T. Chronotypes in the US - Influence of age and sex. PLoS One. 2017;12: e0178782. doi:10.1371/journal.pone.0178782

50. Clark LA, Watson D, Leeka J. Diurnal variation in the positive affects. Motiv Emot. 1989;13: 205–234. doi:10.1007/bf00995536

51. Baglioni C, Spiegelhalder K, Lombardo C, Riemann D. Sleep and emotions: a focus on insomnia. Sleep Med Rev. 2010;14: 227–238. doi:10.1016/j.smrv.2009.10.007

52. Wittmann M, Dinich J, Merrow M, Roenneberg T. Social jetlag: misalignment of biological and social time. Chronobiol Int. 2006;23: 497–509. doi:10.1080/07420520500545979

