## Supplementary Tables for "Time-of-day related fluctuations of self-belief formation"

| *Supplementary Table 1.* Likelihood ratio test of linear mixed-effects models on learning rates across time-of-day intervals and estimates of the final model. | | | | | | |
| --- | --- | --- | --- | --- | --- | --- |
| **Added fixed effects** | **Model fit** | | | **Chi-square test against nested** | | |
|  | AIC | BIC | LL | df | *Χ2* | *p* |
| Intercept | -321.8583 | -309.3120 | 163.9291 |  |  |  |
| Time | -326.0643 | -296.7897 | 170.0321 | 4 | 12.21 | .016* |
| PE-valence | -375.7693 | -342.3126 | 195.8846 | 1 | 51.70 | <.001*** |
| Time*PE-valence | -370.0385 | -319.8535 | 197.0193 | 4 | 2.27 | .686 |

| **Fixed effects** | Beta | *SE* | 95% CI | | *t* | *p* |
| --- | --- | --- | --- | --- | --- | --- |
|  |  |  | lower | upper |  |  |
| Intercept | 0.21 | 0.02 | 0.18 | 0.25 | 12.28 | <.001*** |
| 10:00-12:59am vs. 7:00-9:59am | 0.03 | 0.02 | -0.02 | 0.07 | 1.06 | .29 |
| 1:00-3:59pm vs. 7:00-9:59am | -0.02 | 0.02 | -0.07 | 0.02 | -0.96 | .339 |
| 4:00-6:59pm vs. 7:00-9:59am | 0.00 | 0.03 | -0.06 | 0.06 | -0.01 | .995 |
| 7:00-9:59pm vs. 7:00-9:59am | 0.07 | 0.03 | 0.02 | 0.12 | 2.52 | .012* |
| PE-Valence | 0.10 | 0.01 | 0.07 | 0.12 | 7.58 | <.001*** |
| **Random effects** | | | *Variance* | | *SD* | |
| Participant (Intercept) | | | 0.01 | | 0.08 | |
| **Model fit** | | | Marginal *R^2^* | | Conditional *R^2^* | |
|  | | | 0.11 | | 0.33 | |
| *Note.* AIC = Akaike information criterion; BIC = Bayesian information criterion; LL = log-likelihood; *df* = degrees of freedom; *SE* = standard error of the mean; 95% CI = 95% confidence intervals of beta estimates. Equation of final model (shaded line): learning rate ~ (1\|subject) + time + PE-valence. *p*-values for fixed effects were calculated using Satterthwaite's approach. | | | | | | |

| *Supplementary Table 2.* Likelihood ratio test of linear and polynomial mixed-effects models on learning rates across time-of-day (standardized) and estimates of the final model. | | | | | | |
| --- | --- | --- | --- | --- | --- | --- |
| **Added fixed effects** | **Model fit** | | | **Chi-square test against nested** | | |
|  | AIC | BIC | LL | df | *Χ2* | *p* |
| Linear base model: Time + PE-valence | -371.7720 | -350.8615 | 190.8860 |  |  |  |
| Interaction: Time*PE-valence | -369.9692 | -344.8767 | 190.9846 | 1 | 0.2 | .657 |
| **Interaction term removed; next model tested against linear base model** | | | | | | |
| Linear, quadratic & cubic terms for Time | -376.1657 | -346.8911 | 195.0828 | 2 | 8.39 | .015* |
| Polynomial interactions Time*PE-valence | -371.3190 | -329.4982 | 195.6595 | 3 | 1.15 | .764 |

| **Fixed effects** | Beta | *SE* | 95% CI | | *t* | *p* |
| --- | --- | --- | --- | --- | --- | --- |
|  |  |  | lower | upper |  |  |
| Intercept | 0.22 | 0.01 | 0.19 | 0.24 | 15.08 | <.001*** |
| PE-valence | 0.1 | 0.01 | 0.07 | 0.12 | 7.58 | <.001*** |
| Time (linear) | -0.01 | 5.9e-03 | -0.02 | 7.1e-04 | -1.84 | .067 |
| Time (quadratic) | -5.8e-04 | 9e-04 | -2.3e-03 | 1.2e-03 | -0.64 | .525 |
| Time (cubic) | 4e-04 | 1.9e-04 | 3.4e-05 | 7.6e-04 | 2.14 | .033* |
| **Random effects** | | | *Variance* | | *SD* | |
| Participant (Intercept) | | | 0.01 | | 0.08 | |
| **Model fit** | | | Marginal *R^2^* | | Conditional *R^2^* | |
|  | | | 0.1 | | 0.33 | |
| *Note.* AIC = Akaike information criterion; BIC = Bayesian information criterion; LL = log-likelihood; *df* = degrees of freedom; *SE* = standard error of the mean; 95% CI = 95% confidence intervals of beta estimates. Equation of final model (shaded line): learning rate ~ (1\|subject) + poly(time, 3) + PE-valence.. *p*-values for fixed effects were calculated using Satterthwaite's approach. | | | | | | |
